# Interferon-γ Increases Sensitivity to Chemotherapy and Provides Immunotherapy Targets in Models of Metastatic Castration-Resistant Prostate Cancer

**DOI:** 10.1101/2021.08.20.457028

**Authors:** Dimitrios Korentzelos, Alan Wells, Amanda M. Clark

**Affiliations:** Department of Pathology, University of Pittsburgh, Pittsburgh, PA 15261, USA; UPMC Hillman Cancer Center, University of Pittsburgh, Pittsburgh, PA 15213, USA; VA Pittsburgh Healthcare System, Pittsburgh, PA 15213, USA; Department of Bioengineering, University of Pittsburgh, Pittsburgh, PA 15260, USA; Department of Computational & Systems Biology, University of Pittsburgh, Pittsburgh, PA 15260, USA; Pittsburgh Liver Research Center, University of Pittsburgh and UPMC, Pittsburgh, PA 15261, USA

## Abstract

Interferon-γ (IFNγ) is a cytokine with limited evidence of benefit in cancer clinical trials to date. However, it could potentially play a role in potentiating anti-tumor immunity in the immunologically “cold” metastatic castration-resistant prostate cancer (mCRPC) by inducing antigen presentation pathways and concurrently providing targets for immune checkpoint blockade therapy. Moreover, it could additionally increase sensitivity to chemotherapy based on its pleiotropic effects on cell phenotype. Here, we show that IFNγ treatment induced expression of major histocompatibility class-I (MHC-I) genes and PD-L1 in prostate cancer cells in vitro. Furthermore, IFNγ treatment led to a decrease in E-cadherin expression with a consequent increase in sensitivity to chemotherapy in vitro. In an in vivo murine tumor model of spontaneous metastatic prostate cancer, IFNγ systemic pretreatment upregulated the expression of HLA-A and decreased E-cadherin expression in the primary tumor, and more importantly in the metastatic site led to increased apoptosis and limited micrometastases in combination with paclitaxel treatment compared to diffuse metastatic disease in control and monotherapy treatment groups. These findings suggest that IFNγ may be useful in combinatorial regimens to induce sensitivity to immunotherapy and chemotherapy in hepatic metastases of mCRPC.

## Introduction

Interferon-γ (IFNγ) is a multifunctional cytokine currently approved by the FDA for the treatment of osteopetrosis and chronic granulomatous disease (1). As IFNγ also induces a strong antitumor immune response via T-helper 1 cell polarization, cytotoxic lymphocyte activation, and increased dendritic cell tumoricidal activity (2), it was extensively evaluated as a single agent cancer immunotherapy in multiple clinical trials during the 1990s. However, it was associated with inconsistent results, and as a consequence eventually these efforts were largely abandoned. In the current era of immunotherapy, it is understood that these controversial results are largely due to the dual roles of IFNγ; it is also able to promote tumor development and progression, particularly by upregulating aggression and immune checkpoint molecules, such as the PD-1/PD-L1 axis (3).

Conversely, one of the other major effects of IFNγ is upregulation of major histocompatibility class-I (MHC-I) gene expression and therefore in the setting of cancer, increased tumor antigen processing and presentation leading to improved T-cell recognition and cytotoxicity (4–8). Several studies have shown a strong correlation between markers of antigen presentation and response to immune checkpoint blockers (ICBs) targeting the PD-1/PD-L1 axis (9–12). This would make IFNγ a promising strategy in the case of relatively “immune cold” cancers, such as metastatic castration-resistant prostate cancer (mCRPC), which is characterized by a low tumor mutational burden and few tumor infiltrating T cells, and are relatively resistant to ICBs (13). Indeed, clinical trials of anti-PD-1/PD-L1 monotherapies in prostate cancer have exhibited limited benefit thus far, including a recent phase III randomized clinical trial of atezolizumab failing to meet its primary overall survival (OS) endpoint (14–17). Moreover, with regards to the liver, which represents the second most common metastatic site for prostate cancer after bone and the one with the worst prognosis (18), our current understanding is that its immunologic characteristics are unique in facilitating metastatic expansion via diminished immunity to neoantigens entering through the portal circulation (19).

Metastatic cancers are not only frequently able to avoid immune surveillance, but also show resistance to chemotherapy either in a primary, intrinsic or a secondary, adaptive manner (20). In the case of dormant micrometastases, our group has previously demonstrated that one of the factors conferring survival advantage with regards to avoidance of chemotherapy is E-cadherin re-expression in an autocrine or paracrine fashion (21–23). This is particularly evident in the liver, where hepatocytes promote p38- and ERK-mediated phenotypic switching in mCRPC cells supporting tumor cell survival in the face of death signals (24, 25). This makes IFNγ even more promising as a potential part of a combinatorial treatment strategy in mCRPC, as it has been shown that decreased membranous expression of E-cadherin is evoked by IFNγ in a Fyn Kinase-dependent manner, and additionally, IFNγ-induced IFIT5 suppresses E-cadherin in prostate cancer via altered miRNA processing (26, 27).

Based on the above, in this study we explored the effects of IFNγ in the induction of MHC-I and PD-L1 expression and the suppression of E-cadherin expression in preclinical models of mCRPC, as well as the impact of these effects on sensitivity of mCRPC to chemotherapy.

## Results

### IFNγ induces upregulation of MHC-I and PD-L1 and downregulation of E-cadherin in mCRPC cells

To test the effect of IFNγ on MHC-I, PD-L1 and E-cadherin expression in mCRPC cells, we evaluated one normal prostate cell line (RWPE) as well as different prostate cancer cell lines representing both androgen-dependent (LnCaP) and androgen-independent (DU145, PC3) subtypes of metastatic prostate cancer for their response to IFNγ (5 ng/mL, 48 hour treatment). Furthermore, to achieve a better representation across the spectrum of cancer-associated epithelial-to-mesenchymal transition (cEMT), we specifically used cell line variants that express high and low E-cadherin in the DU-145 (DU-H, DU-L) and PC3 (PC3-H, PC3-L) cell lines. E-cadherin and PD-L1 levels or E-cadherin and MHC-I levels were measured via flow cytometry with and without membrane permeabilization in order to observe total and membranous expression, respectively. Treatment with IFNγ significantly increased both membranous and total expression of MHC-I and PD-L1 in benign RWPE cells as well as DU-H, DU-L, PC3-H and PC3-L cancer cells (Fig. 1A, Fig. S1, S2, S3C, S4C). Interestingly, no differences were observed in LnCaP cells (Fig. S5C) in PD-L1 or MHC-I expression. The aforementioned results were corroborated quantitatively by immunoblotting and qualitatively by immunofluorescence (Fig. 1B-D, Fig. S3A,B, S4A,B, S5A,B).

**Fig. 1.**
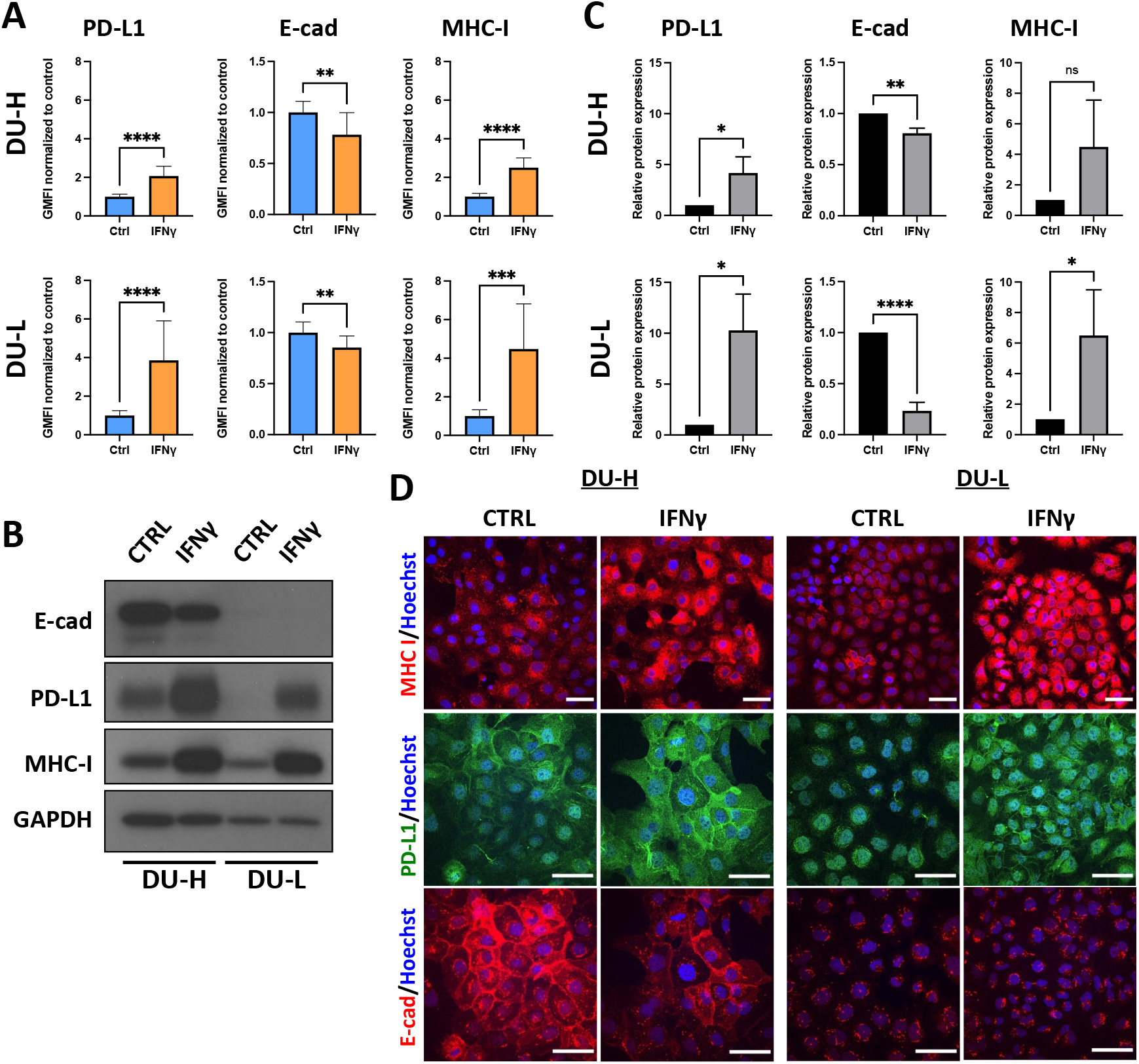
IFNγ treatment alters the expression of MHC-I, PD-L1 and E-cadherin in mCRPC cells. (A) Geometric Mean Fluorescence Intensity (GMFI) of E-cadherin, MHC-I, and PD-L1 total expression in DU-H and DU-L cells after treatment with control or IFNγ (5 ng/mL) for 48 hours, determined by flow cytometry. (B) Immonoblot of E-cadherin, PD-L1, and MHC-I in DU-H and DU-L cells after control or IFNγ (5 ng/mL) treatment for 48 hours, with GAPDH as loading control. (C) Quantification of immunoblots with fold change compared with control. Data shown as mean ± SD of three independent experiments. Student t-test *p<0.05, **p<0.005, ****p<0.0001, ns, not significant. (D) Representative immunofluorescence (IF) images of staining MHC-I (red), PD-L1 (green), E-cadherin (red), and Hoechst 33342 (blue) in DU-H and DU-L. Cells were treated with control or IFNγ (5 ng/mL) for 48 hours. All scale bars, 50 μm. Data shown as mean ± SD of at least three independent experiments. Student t-test, ***, p<0.001.

Regarding changes in E-cadherin expression, IFNγ induced significant downregulation of total E-cadherin in benign RWPE cells and DU-H, DU-L, and PC3-H cancer cells (Fig. 1A, Fig. S1, S2, S3C). On the other hand, there was no change in total E-cadherin expression in PC3-L cells (Fig. S4C), while there was a mild (10%) decrease in LnCaP cells (p=0.10; Fig. S5C). Interestingly, while all cancer cell lines showed evidence of lower membranous E-cadherin expression after IFNγ treatment, statistical significance was only achieved for PC3-H, DU-L and DU-H cells (Fig. S2, Fig. S3C) and no change was noticed in RWPE cells (Fig. S1). Strikingly, the decrease in membranous E-cadherin in PC3-H cells following IFNγ was almost 39% (p=0.0007; Fig. S3C). Again, immunofluorescence and immunoblots further confirmed the aforementioned findings (Fig. 1B-D, Fig. S3A,B, S4A,B, S5A,B).

### IFNγ potentiates response to chemotherapy in mCRPC in vitro

As stated above, it has been previously shown by our group that hepatocyte-induced E-cadherin re-expression in breast and prostate cancer cells leads to increased chemoresistance (21). Thus, we next asked whether IFNγ-induced E-cadherin downregulation could effectively sensitize mCRPC cells to chemotherapy. Concordantly, we treated DU-H, DU-L, PC3-H, and PC3-L with IFNγ (5 ng/mL, 48 hour treatment) prior to administration of combination treatment with 1 μM camptothecin (CPT) and 100 ng/mL tumor necrosis factor-related apoptosis inducing ligand (TRAIL). This combination of a cytotoxic chemotherapeutic agent with a death cytokine was selected because it is physiologically relevant as dormant prostate cancer metastases are resistant to death inducing signals, whether from chemotherapies or cytokines. Moreover, our group has previously shown that the aforementioned cell lines were more sensitive to concurrent treatment than monotherapy with any of these agents (25). Cleaved caspase 3 was used as a surrogate marker for apoptotic activity and membranous E-cadherin was measured as previously described (25). In all cases, cleaved caspase 3 was significantly increased in the IFNγ-pretreatment mCRPC cell lines illustrating increased sensitivity to chemotherapy after sensitization with IFNγ (Fig 2). The most striking increase was observed in DU-L cells (164%, p<0.0001) and then in sequential order in PC3-L (74.3%, p=0.0003), PC3-H (73.3%, p<0.0001) and DU-H (13.3%, p=0.03). With regards to membranous E-cadherin expression in IFNγ-treated groups, PC3-L cells demonstrated a reduction by 43% (p=0.005), DU-L and PC3-H cells showed small decreases which did not reach statistical significance, while DU-H showed a small, not significant increase (Fig. 2).

**Fig. 2.**
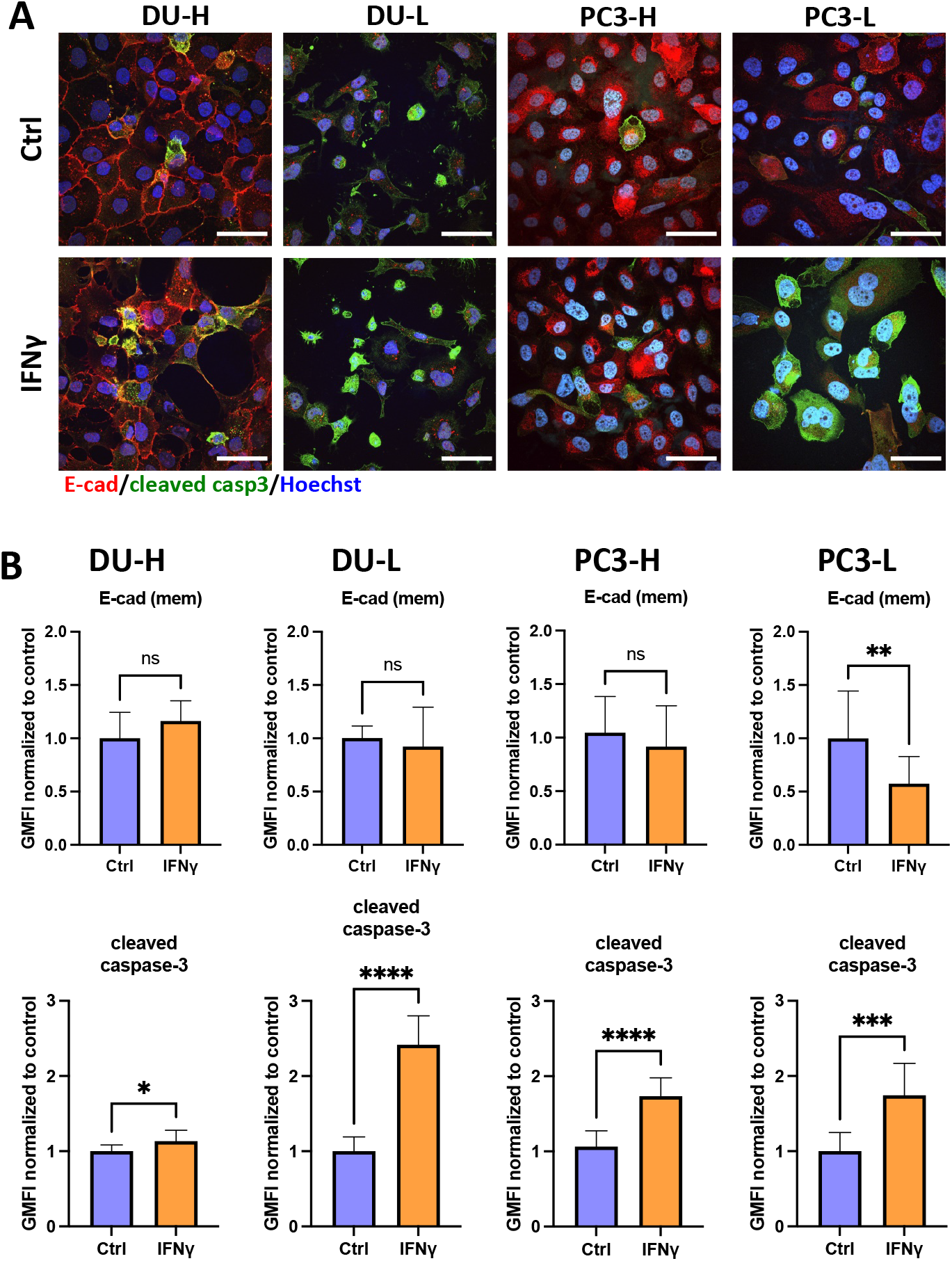
IFNγ influences the response of mCRPC cells to chemotherapy in vitro. (A) Representative IF images of co-staining E-cadherin (red), cleaved caspase 3 (green), and nucleus (Hoechst 33342, blue) in DU-H, DU-L, PC3-H, and PC3-L. Cells were treated with 1 μM camptothecin and 100 ng/ml TRAIL (CPT-TRAIL) for 4 hours after 48 hours of control or IFNγ (5 ng/mL) treatment. All scale bars, 50 μm. (B) GMFI of cleaved caspase 3 and membranous E-cadherin expression in DU-H, DU-L, PC3-H and PC3-L cells determined by flow cytometry. Data shown as mean ± SD. Student t-test, *p<0.05, **p<0.005, ***, p<0.001, ****p<0.0001.

### IFNγ decreases E-cadherin expression and potentiates response to chemotherapy in mCRPC in vivo

The next step was to address whether IFNγ-induced E-cadherin downregulation renders mCRPC more sensitive to chemotherapy in a preclinical murine model of spontaneous mCRPC liver metastasis. We injected DU-L cells into the spleen of mice, as our group has previously shown this to be a valid model to consistently produce liver metastases by day 18 post-injection (25). Mice were divided into four treatment groups: control, single IFNγ dose on day 18, chemotherapy, and combination of IFNγ on day 18 and chemotherapy (Fig. 3A). The drug used was a taxane (paclitaxel, PAC), a class of chemotherapy suggested by the National Comprehensive Cancer Network for patients with advanced prostate cancer. More specifically, paclitaxel was chosen based on prior literature on mice for dosing and also prior work from our group. Mouse weight was recorded prior to each drug injection to monitor for potential side effects; no obvious weight loss (>5%) was observed in any of the mice. The mice were euthanized on day 30 after five rounds of PAC treatment.

**Fig. 3.**
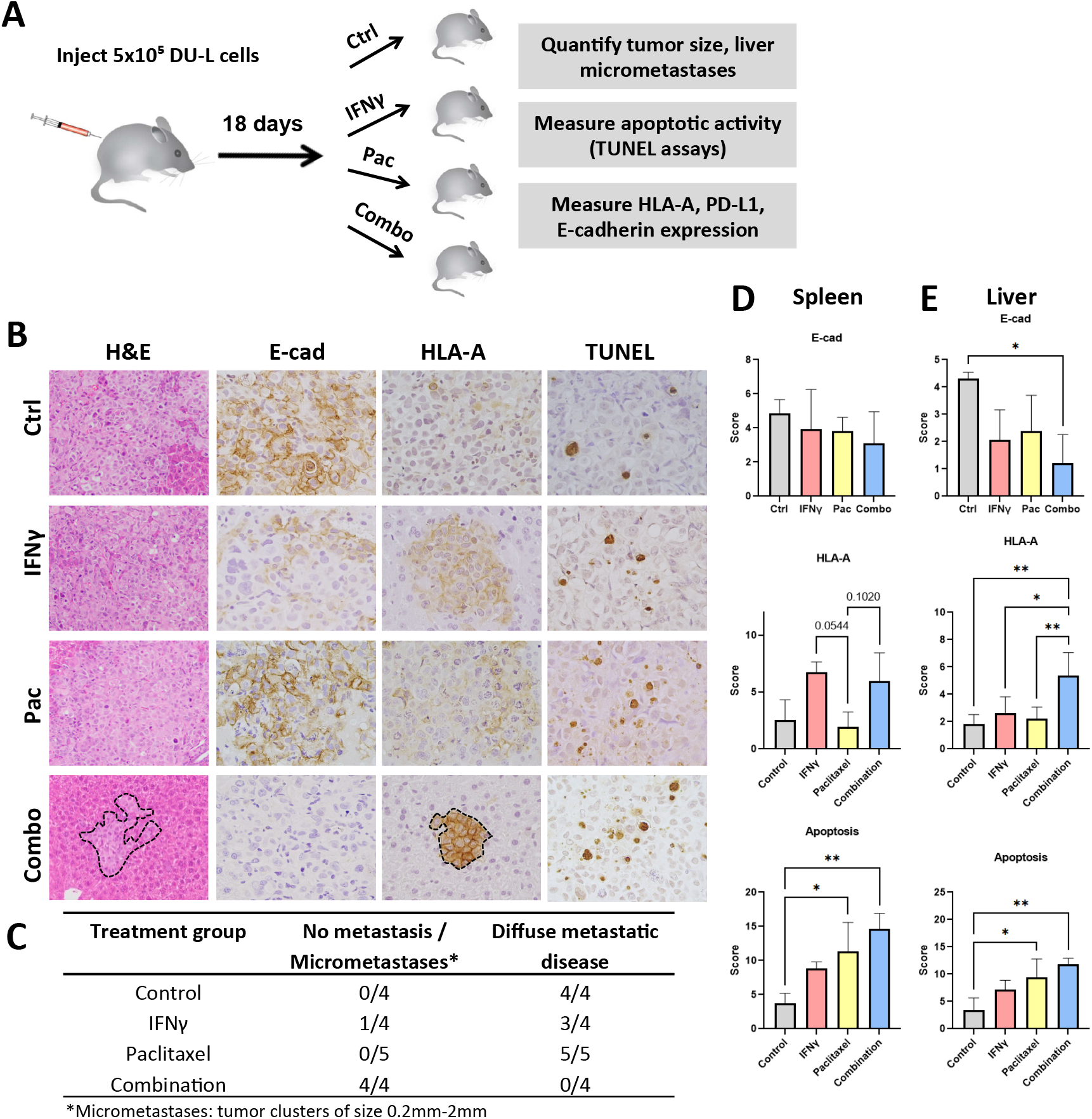
Alterations in HLA-A and E-cadherin expression and chemosensitivity of mCRPC after IFNγ pretreatment in vivo. (A) Experimental outline to test the effect of IFNγ pretreatment on HLA-A and E-cadherin expression, as well as sensitivity to chemotherapy in DU-L cells in vivo. (B) Limited liver metastatic tumor growth of DU-L prostate cancer cells in IFNγ and paclitaxel combination group. H&E images at 200x magnification and all others at 400x magnification. Tumor is not outlined where the vast majority of the captured image is tumor. (C) Enumeration of liver metastasis in each mouse. (D) Representative H&E, HLA-A, E-cadherin and TUNEL staining in the metastatic tumors at completion of the study. (E) Quantification of HLA-A and E-cadherin expression, and TUNEL assays. Data shown as mean ± SD. Analysis of variance (ANOVA) or Kruskal-Wallis with Dunn’s multiple comparisons test after determination of normality based on Shapiro-Wilk test, *p<0.05, **p<0.005, ***, p<0.001, ****p<0.0001.

All the primary tumors that arose upon intrasplenic injection produced hepatic metastases with the exception of one tumor treated with combination therapy, which did not lead to any detectable metastasis to the liver. Strikingly, all samples in control and PAC treatment groups, and 3/4 samples from IFNγ group, demonstrated a diffuse pattern of liver involvement with almost complete replacement of the organ by metastatic tumor (Fig 3B,C). One sample from IFNγ group had only a small focus of micrometastasis, but also a well-circumscribed tumor nodule in the spleen, in contrast to the other mice that had diffuse infiltration of the spleen by tumor. Nevertheless, the 3 samples from the combination group that showed metastatic deposits, either had multiple small, well-circumscribed metastatic foci (2/4 samples) occupying less than 10% of the organ, or only rare micrometastases (1/4 samples). This difference in diffuse metastasis versus no metastasis/micrometastases between combination and all other groups combined was statistically significant (chi square test, χ^2^=13.39, df=3, p=0.0039). Consistently, E-cadherin expression was decreased in the combination group compared to control, while interestingly a trend was not only observed in the IFNγ group as expected but also in the PAC group as well (Fig 3B, E). Nevertheless, this decrease only reached the level of statistical significance in the liver and in the comparison between control and combinatorial treatment (Fig. 3E). This result clearly demonstrates that the pretreatment with IFNγ leads to a decrease in E-cadherin expression and sensitizes the liver metastases of mCRPC to chemotherapy.

We then pursued further mechanistic insight into the increased chemosensitivity of mCRPC in response to IFNγ by performing TUNEL assays to evaluate apoptosis. Apoptotic activity was significantly increased in PAC and combination groups compared to control and IFNγ, albeit there was no difference between PAC and combination groups (Fig. 3E).

### Effects of IFNγ on MHC-I levels in mCRPC in vivo

To confirm that treatment with IFNγ induces upregulation of PD-L1 and MHC-I in liver metastases of mCRPC we measured the levels of the two markers by IHC (Fig. 3B,D,E). With regards to HLA-A, both IFNγ and combination treatment groups showed increased expression in the primary tumors in the spleen, although did not reach statistical significance (Fig 3B,D). Interestingly, the combination group demonstrated a significant increase in HLA-A in the liver metastases and furthermore, the difference between IFNγ and combination groups was also significant (Fig 3E). The pattern of staining was predominantly membranous, although some cytoplasmic staining could also be appreciated, a finding consistent with the extensive posttranslational modifications of the protein occurring predominantly in the Golgi apparatus (28). Unfortunately, unlike in vitro findings, immunohistochemical staining with PD-L1 was minimal-to-absent in all four treatment groups, with a non-specific speckled pattern rather than membranous when present, both in the primary and in the metastatic setting, despite the use of a clinically validated antibody at a high concentration, prohibiting any further accurate analyses (Fig. S6).

## Discussion

Herein, we explored the effects of IFNγ to promote the expression of MHC-I and PD-L1 in mCRPC preclinical models, as well as its potential to downregulate E-cadherin in order to render mCRPC more sensitive to chemotherapy. Among multiple prostate cancer cell lines, spanning diverse molecular subtypes both in terms of AR expression as well as position along the cEMT spectrum, we found that IFNγ upregulated MHC-I expression both in vitro and in vivo, and PD-L1 expression was upregulated in vitro (corroboration in vivo was not technically possible). Moreover, E-cadherin expression was reduced in response to IFNγ pretreatment, and this led to increased apoptosis, as measured based on cleaved caspase 3 levels, upon exposure of mCRPC cells to a combination of a chemotherapeutic drug with a death cytokine. Strikingly, the combination of IFNγ with subsequent administration of chemotherapy (paclitaxel) led to limited metastatic disease (micrometastases) in vivo compared to the diffuse infiltration of the liver by metastatic tumors in the case of control and monotherapy with either IFNγ or paclitaxel. The present work is a novel approach and makes it clear that IFNγ has potential as an agent that may provide targets for immunotherapy and concurrently increase the sensitivity of mCRPC to chemotherapy.

The study has a number of limitations and open questions for further study. The paradoxical mild and not statistically significant increase in membranous E-cadherin in DU-H cells after IFNγ treatment in the chemosensitivity assays could potentially be explained by the fact that mCRPC cells with lower expression of E-cadherin may be more easily damaged by chemotherapy. Another limitation was the lack of PD-L1 staining in spleens and livers of mice from all groups despite the use of a clinically validated antibody. Nevertheless, this can be explained by the very rare/low expression of PD-L1 in mCRPC (29), which means that even after IFNγ-induced PD-L1 upregulation, the PD-L1 levels might still fall under the analytical sensitivity of the IHC method. Alternatively, N-linked glycosylation of PD-L1 might potentially hinder its recognition by the PD-L1 antibody used, which means that a deglycosylation step would be required in order to prevent a false-negative result (30).

Although interest in IFNγ has waned over the last years in lieu of targeted agents and because of its confounding pro- and anti-tumoral actions, the failure of ICBs in the case of immune cold tumors such as mCRPC put it again into consideration. Recently, Zhang et al reported their results from a phase 0 clinical trial of systemic IFNγ in two other immune cold tumors, synovial sarcoma and myxoid/round cell liposarcoma, where IFNγ resulted in increased MHC-I expression and significant T-cell infiltration with better tumor antigen presentation and less exhausted phenotypes of the TILs (1). Furthermore, the IFNγ-induced PD-L1 upregulation which was traditionally considered a major player driving pro-tumoral effects of IFNγ, could potentially now represent the basis for powerful combinatorial approaches with ICBs, a phenomenon similar to the positive predictive value of an otherwise negative prognostic biomarker, HER2/neu amplification, in breast cancer. However, these in vitro and in vivo findings have to be tested in further preclinical studies and future clinical trials with particular attention to the optimal timing of IFNγ administration before the intervention with a PD-1/PD-L1 inhibitor.

## Methods

### Cell lines

DU145 (both variants herein termed DU-L [E-cadherin low expressing] and DU-H [E-cadherin high expressing]), PC3 (both variants herein termed PC3-L [E-cadherin low expressing] and PC3-H [E-cadherin high expressing]) and LnCaP human prostate cancer cell lines, and RWPE-1 normal prostate cell line, were purchased from the American Type Culture Collection (Manassas, VA, USA). DU-H/DU-L and PC3-H/PC3-L cells were maintained in DMEM and F-12K media, respectively. LnCaP cells were maintained in RPMI 1640 media and RWPE-1 cells in Keratinocyte Serum Free Medium (K-SFM). Cells were seeded into 6-well plates and after 24 hours treated with IFNγ (5 ng/mL) or control (PBS) for 48 hours. Cells were harvested for immunoblot or flow cytometry, or fixed for immunofluorescence.

### Immunofluorescence

Cells on coverslips were fixed in cold 2% paraformaldehyde in PBS, and then permeabilized in 0.2% Triton X-100 in PBS for 20 minutes. Cells were subsequently washed in PBS, blocked for 1 hour at room temperature (RT) in PBS containing 2% bovine serum albumin (Sigma-Aldrich). Fixed cells were then incubated with the primary antibody in 2% BSA/PBS solution overnight at 4°C. Cells were washed in PBS three times, and the secondary antibody was added in 1% BSA/PBS solution for 1 hour at RT in the dark. Cells were washed three times in PBS and then incubated for 2 minutes in Hoechst 33342 solution, after which they were washed in PBS and mounted. Confocal images were obtained on an Olympus upright Fluoview 2000 confocal microscope (Center for Biologic Imaging, University of Pittsburgh, supported by NIH #1S10OD019973-01) using a 60x (UPlanApo NA=1.42) or 20x (UPlanSApo NA=0.85) objective. The primary antibodies used were rabbit anti-human cleaved caspase-3 (9661, Cell Signaling Technology), mouse anti-human E-cadherin (Clone 36, BD), mouse antihuman HLA Class 1 ABC antibody (EMR8-5, Abcam) and mouse anti-human PD-L1 (28-8, Abcam). Secondary antibodies used were goat anti-mouse Alexa Fluor^®^ 488 or goat anti-rabbit Alexa Fluor^®^ 647 (Life Technologies). Hoechst 33342 was applied for nuclei counterstaining.

### Immunoblot

Whole-cell extracts were prepared by lysing cells for 15 minutes on ice in RIPA lysis buffer (50mM Tris-HCl (pH 7.5), 150mM NaCl, 1.0% NP-40, 0.1% SDS, and 0.1% Na-deoxycholic acid) supplemented with protease cocktail inhibitor (Thermo Fisher). Cellular lysates were assayed for protein concentration using Pierce™ BCA Protein Assay Kit in 96-well plates using a microplate reader. Whole cell lysates were separated through 7.5% SDS-polyacrylamide gels and transferred to nitrocellulose membrane (Bio-Rad Laboratories, Inc.). Membranes were blocked with 5% milk powder in 0.1% Tween 20 in 1x Tris-Buffered Saline (TBST) for 1 hour at RT followed by incubation with primary antibodies diluted in 5% milk/TBST. Pierce ECL Western blotting substrate (Thermo Scientific) was used to visualize protein levels with light sensitive-films (Thermo Scientific CL-XPosure Film). Immunoblots were quantified using ImageJ software. The following antibodies were used in this study: E-cadherin (24E10, Cell Signaling Technology and clone 36, BD), PD-L1 (E1L3N; Cell Signaling Technology), HLA Class 1 ABC (EMR8-5, Abcam), and GAPDH (D16H11; Cell Signaling Technology).

### Flow cytometry

Cells were rinsed with warm PBS and harvested with a non-enzyme cell dissociation buffer (Life Technology). After centrifugation, cells were fixed with 2% paraformaldehyde in PBS for 30 minutes then rinsed with 1% FBS/PBS. For total expression, cells were permeabilized with 0.2% Triton X-100 in PBS for 20 minutes and subsequently washed in PBS. Both membranous and total expression samples were then incubated with an FcR Blocker along with the primary antibodies in 1% FBS/PBS for 30 minutes at 4 °C. They were then washed with 1% FBS/PBS and incubated with secondary antibodies for 30 minutes at 4 °C. The primary antibodies used were rabbit anti-human cleaved caspase-3 (9661, Cell Signaling Technology), mouse anti-human E-cadherin (Clone 36, BD), mouse anti-human HLA Class 1 ABC antibody (EMR8-5, Abcam) and mouse anti-human PD-L1 (28-8, Abcam). Secondary antibodies used were goat anti-mouse Alexa Fluor^®^ 488 or goat anti-rabbit Alexa Fluor^®^ 647 (Life Technologies). Cells were sorted on BD FACSCalibur™ and analyzed with FlowJo (v10.6.2).

### Chemoresistance assay

DU145 and PC3 cells were treated with 5 ng/mL IFNγ or control for 48 hours and then serum starved overnight. Subsequently, cells were treated with a combination of 1 *μ*M camptothecin (Sigma-Aldrich) and 10 ng/mL recombinant human TRAIL (Life Technologies, PH1634) in serum-free medium for 4 hours, prior to harvesting for flow cytometry. Staining for cleaved caspase-3 and membranous E-cadherin was performed as previously described (25).

### Mouse model of liver metastasis

Animal studies were conducted according to a protocol approved by The Association for Assessment and Accreditation of Laboratory Animal Care-accredited Institutional Animal Care and Use Committees of the Veteran’s Administration Pittsburgh Health System. Seven-week old NOD/SCID gamma mice (00557, The Jackson Laboratory, Bar Harbor, ME) were used. After anesthesia with ketamine/xylazine, pain suppression with long-acting buprenorphine, and sterile surgical exposure of the spleen, half a million of DU145-L cells were injected into the spleen using a 27-gauge needle. The omentum was closed with a running stitch of absorbable suture and the skin wound with metal wound clips. At 2.5 weeks post-injection animals were randomly distributed into four groups: control (n=4), IFNγ (n=4), paclitaxel (n=4) or IFNγ+paclitaxel (n=4). At that point 0.005 μg/kg IFNγ was administered as a single dose in IFNγ and IFNγ+paclitaxel groups only. Paclitaxel (Fresenius Kabi, Lake Zurich, IL) was administered at 10 mg/kg body weight by i.p. every 2 days for a total 5 rounds. Mice were monitored for overall health. After 5 weeks the mice were euthanized using a carbon dioxide chamber consistent with AVMA Guidelines on Euthanasia.

### Immunohistochemistry

Tumor sections were embedded in paraffin, and thin histologic sections (4-5 μm) were prepared and stained with hematoxylin and eosin following standard protocols (performed by the Pitt Biospecimen Core; P30CA047904). For all other stains, tissue sections were deparaffinized in xylene and rehydrated in ethanol following treatment in preheated target retrieval solution. Following washes, serum-free blocking solution was applied for 30 minutes at RT. Expression of E-cadherin, HLA-A, and PD-L1 was determined using rabbit monoclonal antibodies: E-cadherin (24E10; Cell Signaling Technology; 1:100), HLA-A (EP1395Y; Abcam; 1:200) and PD-L1 (28-8; Abcam; 1:100) in formalin fixed, paraffin-embedded tissue sections. The slides were counterstained with hematoxylin, dried and mounted with Permount. Micrographs of the morphology and expression of the markers were captured using the Olympus cellSens software (all at a magnification of x400).

Slides were evaluated by an experienced pathologist. Positively stained tumor cells for E-cadherin or HLA-A were counted in at least five representative high power fields (HPF) for each tumor section. E-cad and HLA-A expression in tumor cells were analyzed using a membranous/cytoplasmic staining algorithm. The staining intensity was scored as 0 (no staining), 1 (weak staining), 2 (moderate staining), or 3 (strong staining) while extension (percentage) of expression were determined as 1 (<10% cells), 2 (10-50% cells) or 3 (>50% cells). The final scores for tumor tissues were determined by multiplying the staining intensity and reactivity extension values (range, 0–9).

### Apoptosis assay

Terminal deoxynucleotidyl transferase dUTP nick end labeling (TUNEL) staining was performed using the ApopTag^®^ Peroxidase in situ Apoptosis Detection Kit (EMD Millipore) according to the manufacturer’s instructions. The number of TUNEL-positive cells was counted in a minimum of five high power fields per slide (400x magnification).

### Statistical analysis

The data in the bar graphs indicate mean ± SD of fold changes in relation to control groups (at least three independent experiments). Statistical analyses were performed using GraphPad Prism 9 (GraphPad Software, Inc., CA, US). The Shapiro-Wilk test was used to test normal distribution of datasets. Two-sided Student *t*-test was used to compare the difference between two independent groups with a parametric data distribution. All of the results containing more than two conditions with a parametric data distribution were analyzed by analysis of variance (ANOVA) post-hoc test (with Tukey’s test to compare all pairs of conditions). Mann-Whitney U test was used to compare the difference between two independent groups with a nonparametric data distribution. Kruskal-Wallis test with post hoc Dunn’s test was applied for multiple comparisons among groups with a nonparametric data distribution. A chisquare test analysis was performed to analyze whether distributions of categorical variables differ from each other. Differences were considered to be statistically significant when the *p* value was below 0.05.

## Acknowledgments

The authors thank members of the Wells and Clark laboratories for their informed suggestions and commentaries.

## Funding

Support for this research was provided by the VA Merit Award program, and the National Institutes of Health (UH3TR000496, GM69668, and GM63569). The funders had no input over any aspects of this work.

## Author contributions

D.K., A.M.C., and A.W. conceived the study, and contributed to the scientific hypotheses, experimental design, methodology, and data interpretation. D.K. and A.M.C. performed experimental work. D.K. wrote the manuscript. A.M.C. and A.W. reviewed and edited the article.

## Competing interests

There are no conflicts of interest to declare.

## Data and materials availability

The data generated for this study are available upon request from the corresponding authors.

## Notes

### Competing Interest Statement

The authors have declared no competing interest.

