## Supplemental Information for "Interferon-γ Increases Sensitivity to Chemotherapy and Provides Immunotherapy Targets in Models of Metastatic Castration-Resistant Prostate Cancer"

**Figures:**

Fig S1. IFNγ induces upregulation of MHC-I and PD-L1, and downregulation of E-cadherin in benign prostate epithelial cells.
Fig S2. Effects of IFNγ on membranous MHC-I, PD-L1, and E-cadherin expression in DU145 cells.
Fig S3. IFNγ induces upregulation of MHC-I and PD-L1, and downregulation of E-cadherin in PC3-H cells.
Fig S4. IFNγ induces upregulation of MHC-I and PD-L1, and downregulation of E-cadherin in PC3-L cells.
Fig S5. Effects of IFNγ on MHC-I, PD-L1 and E-cadherin expression in LnCaP cells.
Fig S6. Effects on HLA-A, PD-L1 and E-cadherin expression and induction of chemosensitivity after IFNγ pretreatment in vivo.

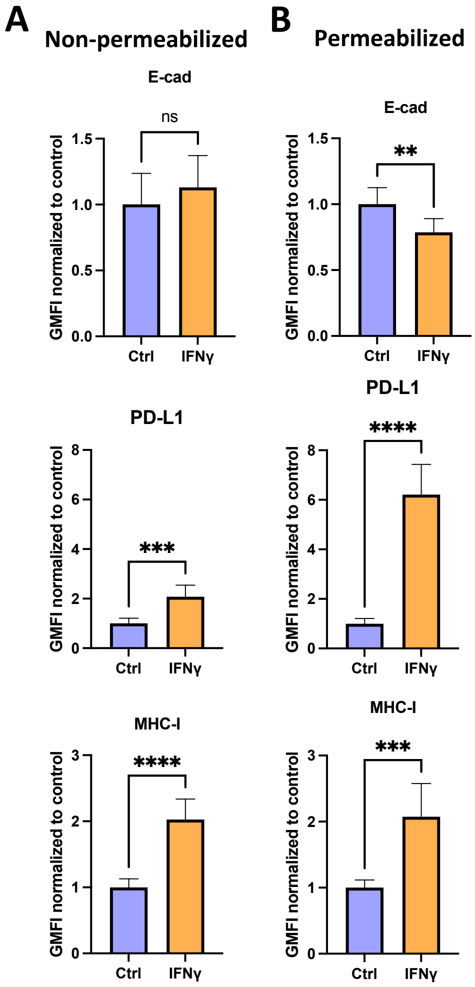


**Fig. S1. IFNγ induces upregulation of MHC-I and PD-L1, and downregulation of E-cadherin in benign prostate epithelial cells.** Geometric Mean Fluorescence Intensity (GMFI) of E-cadherin, MHC-I, and PD-L1 membranous (non-permeabilized cells) and total (permeabilized cells) expression in RWPE cells after treatment with control or IFNγ (5 ng/mL) for 48 hours, determined by flow cytometry. Data shown as mean ± SD. Student t-test, ns, not significant, **p<0.005, ***p<0.001, ****p<0.0001.

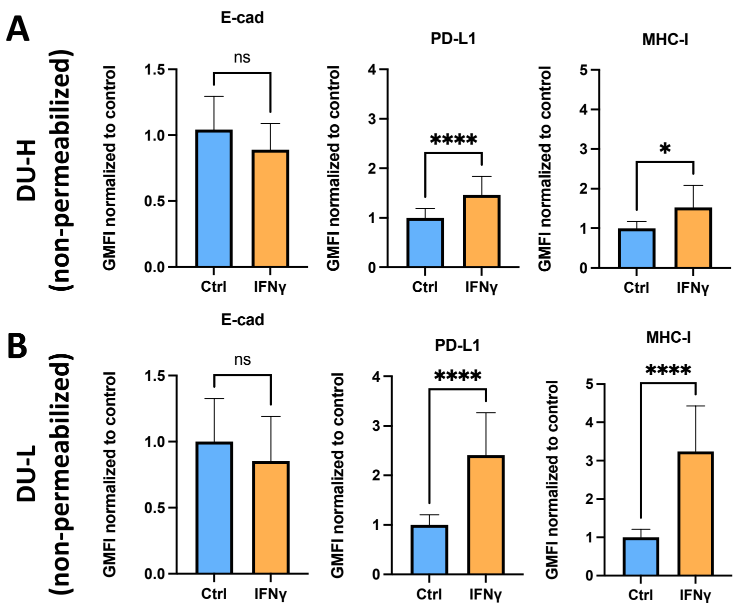


**Fig. S2. Effects of IFNγ on membranous MHC-I, PD-L1 and E-cadherin expression in DU145 cells.** GMFI of membranous E-cadherin, MHC-I, and PD-L1 expression in DU-H and DU-L (non-permeabilized) prostate cancer cells after treatment with control or IFNγ (5 ng/mL) for 48 hours, determined by flow cytometry. Data shown as mean ± SD of at least three independent experiments. Student t-test, ns, not significant, *p<0.05, ****p<0.0001.

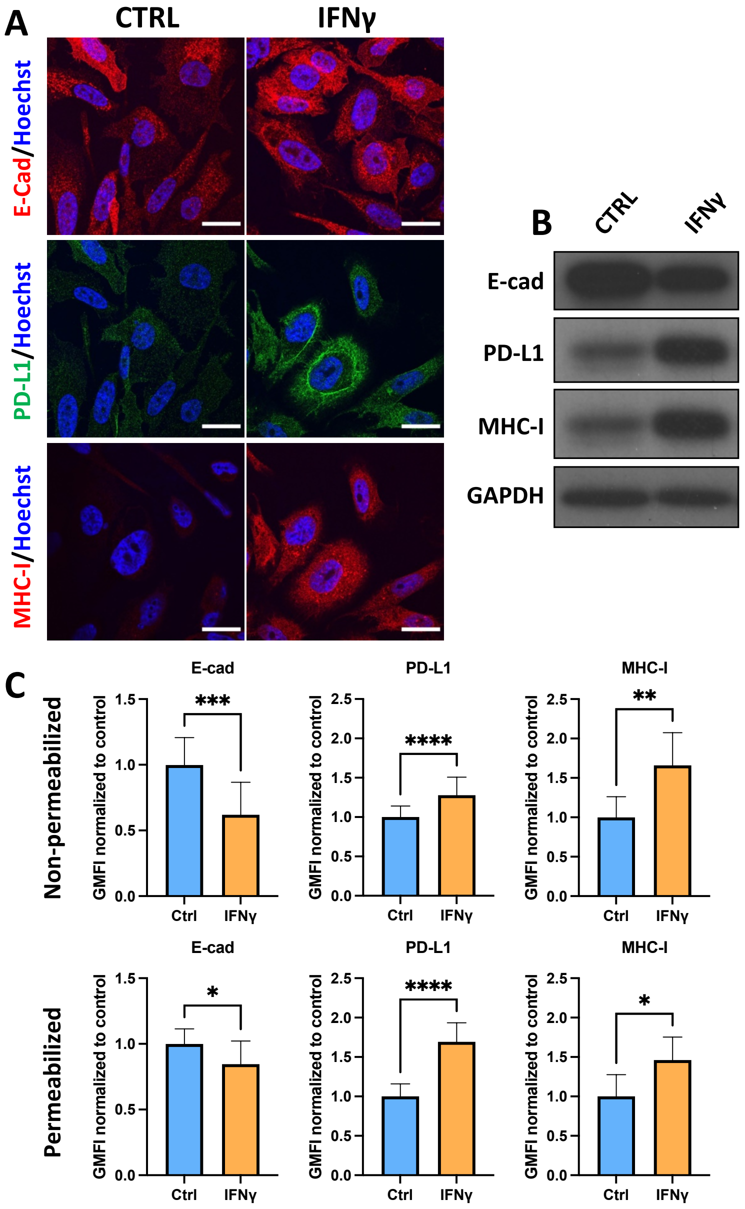


**Fig. S3. IFNγ induces upregulation of MHC-I and PD-L1, and downregulation of E-cadherin in PC3-H cells.** (A) Representative immunofluorescence images of staining MHC-I (red), PD-L1 (green), E-cadherin (red), and Hoechst 33342 (blue) in PC3-H cells. Cells were treated with control or IFNγ (5 ng/mL) for 48 hours. All scale bars, 50 µm. (B) Western blot of E-cadherin, PD-L1, and MHC-I in DU-H and DU-L cells after control or IFNγ (5 ng/mL) treatment for 48 hours, with GAPDH as loading control. (C) GMFI of E-cadherin, MHC-I, and PD-L1 membranous (non-permeabilized cells) and total (permeabilized cells) expression in PC3-H cells after treatment with control or IFNγ (5 ng/mL) for 48 hours, determined by flow cytometry. Data shown as mean ± SD of at least three independent experiments. Student t-test, *$p<0.05$, **$p<0.005$, ***, $p<0.001$, ****$p<0.0001$.

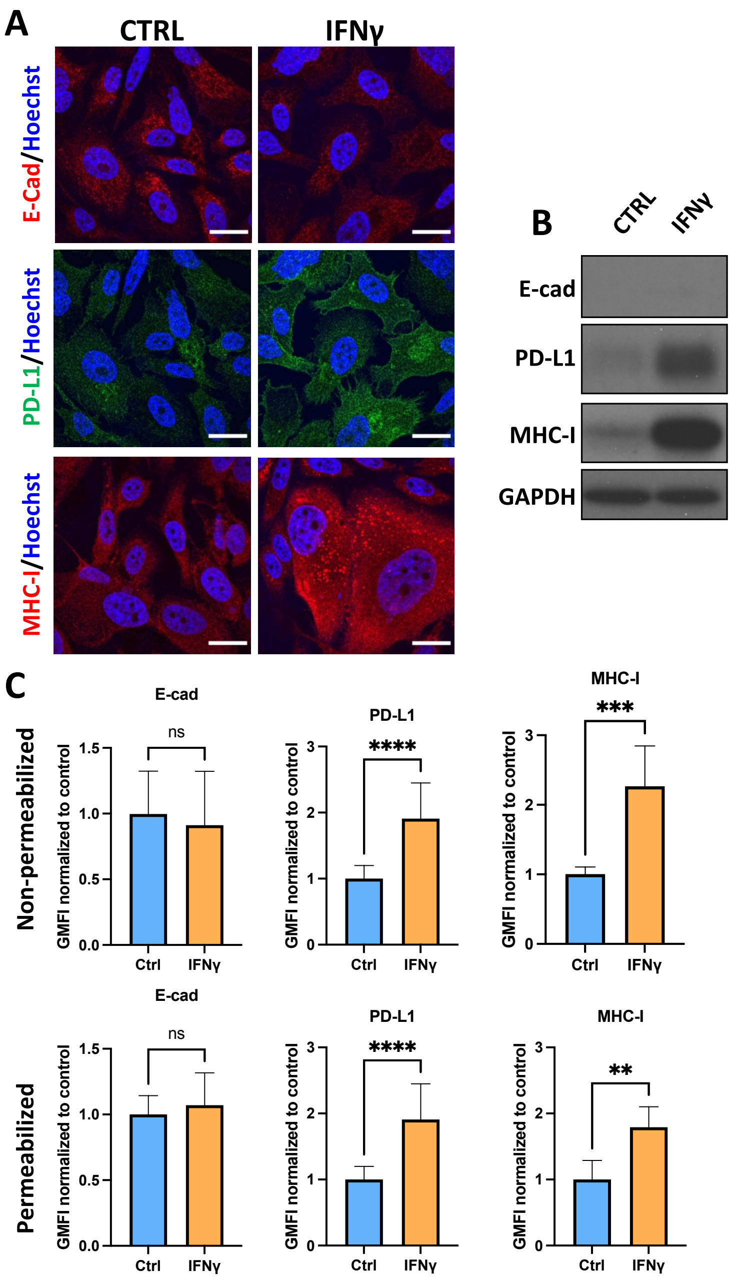


**Fig. S4. IFNγ induces upregulation of MHC-I and PD-L1, and downregulation of E-cadherin in PC3-L cells.** (A) Representative immunofluorescence images of staining MHC-I (red), PD-L1 (green), E-cadherin (red), and Hoechst 33342 (blue) in PC3-L cells. Cells were treated with control or IFNγ (5 ng/mL) for 48 hours. All scale bars, 50 µm. (B) Western blot of E-cadherin, PD-L1, and MHC-I in DU-H and DU-L cells after control or IFNγ (5 ng/mL) treatment for 48 hours, with GAPDH as loading control. (C) GMFI of E-cadherin, MHC-I, and PD-L1 membranous (non-permeabilized cells) and total (permeabilized cells) expression in PC3-L cells after treatment with control or IFNγ (5 ng/mL) for 48 hours, determined by flow cytometry. Data shown as mean ± SD of at least three independent experiments. Student t-test, ns, not significant, $^{**}p<0.005$, ***, $p<0.001$, $^{****}p<0.0001$.

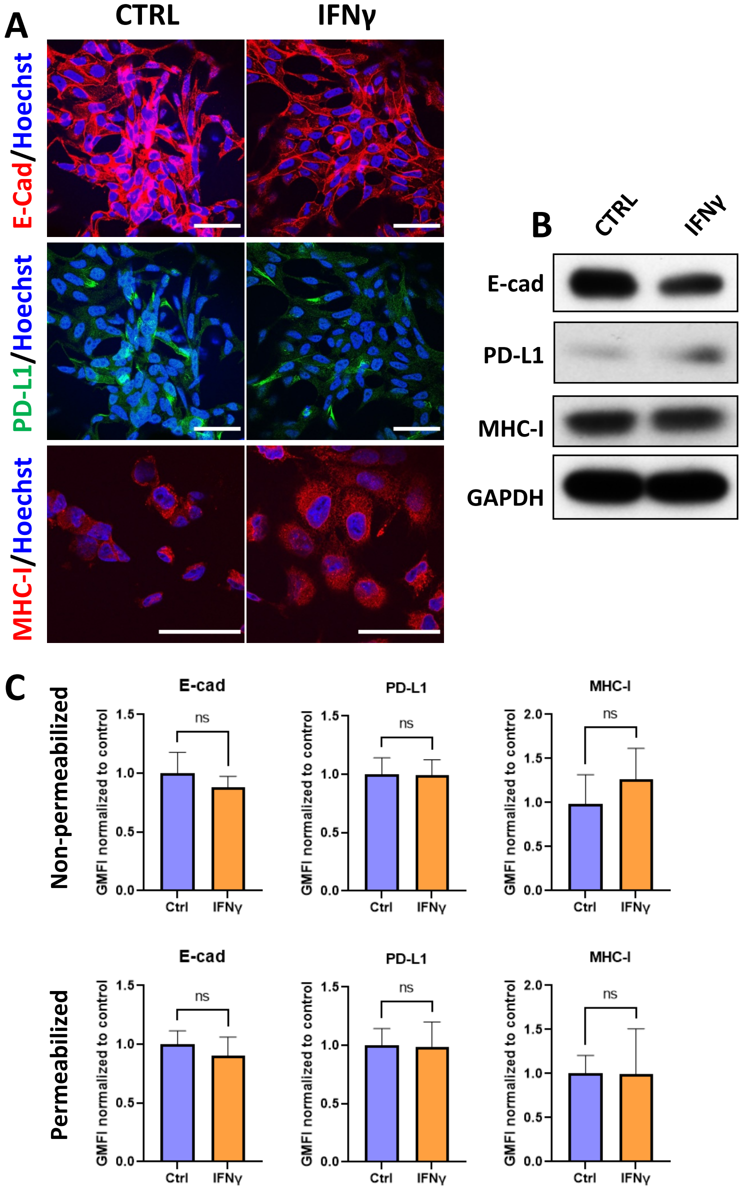


**Fig. S5. Effects of IFNγ on MHC-I, PD-L1 and E-cadherin expression in LnCaP cells.** (A) Representative immunofluorescence images of staining MHC-I (red), PD-L1 (green), E-cadherin (red), and Hoechst 33342 (blue) in LnCaP cells. Cells were treated with control or IFNγ (5 ng/mL) for 48 hours. All scale bars, 50 µm. (B) Western blot of E-cadherin, PD-L1, and MHC-I after control or IFNγ (5 ng/mL) treatment for 48 hours, with GAPDH as loading control. (C) GMFI of E-cadherin, MHC-I, and PD-L1 membranous (non-permeabilized) and total (permeabilized) expression in LnCaP cells after treatment with control or IFNγ (5 ng/mL) for 48 hours, determined by flow cytometry. Data shown as mean±SD of three independent experiments. Student t-test, ns, not significant.

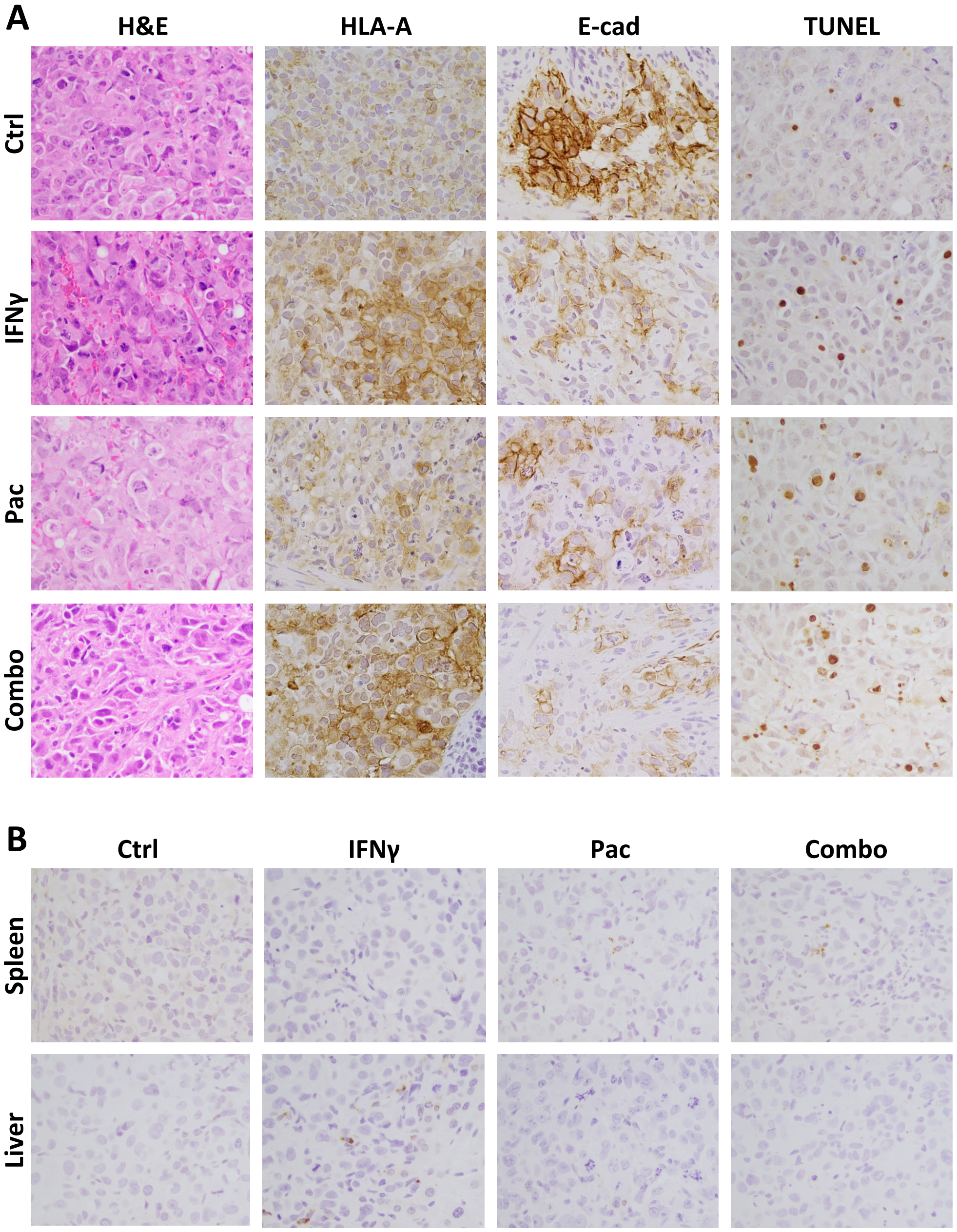


**Fig. S6. Effects on HLA-A, PD-L1 and E-cadherin expression and induction of chemosensitivity after IFNγ pretreatment in vivo.** (A) Representative H&E, HLA-A, E-cadherin and TUNEL staining in the primary (splenic) tumors at completion of the study. (B) Representative PD-L1 staining in primary (splenic) and metastatic (liver) setting at completion of the study. All images at 400x magnification. Tumor is not outlined where the vast majority of the captured image is tumor.
